# Gag proteins encoded by endogenous retroviruses are required for zebrafish development

**DOI:** 10.1101/2024.03.25.586437

**Authors:** Ni-Chen Chang, Jonathan N. Wells, Andrew Y. Wang, Phillip Schofield, Yi-Chia Huang, Vinh H. Truong, Marcos Simoes-Costa, Cédric Feschotte

## Abstract

Transposable elements (TEs) make up the bulk of eukaryotic genomes and examples abound of TE-derived sequences repurposed for organismal function. The process by which TEs become coopted remains obscure because most cases involve ancient, transpositionally inactive elements. Reports of active TEs serving beneficial functions are scarce and often contentious due to difficulties in manipulating repetitive sequences. Here we show that recently active TEs in zebrafish encode products critical for embryonic development. Knockdown and rescue experiments demonstrate that the endogenous retrovirus family BHIKHARI-1 (Bik-1) encodes a Gag protein essential for mesoderm development. Mechanistically, Bik-1 Gag associates with the cell membrane and its ectopic expression in chicken embryos alters cell migration. Similarly, depletion of BHIKHARI-2 Gag, a relative of Bik-1, causes defects in neural crest development in zebrafish. We propose an “addiction” model to explain how active TEs can be integrated into conserved developmental processes.

## Introduction

Transposable elements (TEs) are ubiquitous components of genomes, comprising between 5% and 90% of eukaryotic genomes (*1*, *2*). The mobilization of TEs is a well-documented source of insertional mutagenesis and genome instability (*1*, *3*), and transpositionally active TEs are generally viewed as deleterious to their host organism (*4*, *5*). For the past 50 years, the prevailing dogma has been that TEs are selfish genetic parasites whose evolutionary persistence can be solely attributed to their ability to replicate independently from their host genome (*4*, *6*, *7*). It has also become widely recognized that, once immobilized, TE coding and noncoding sequences can be repurposed into genes and regulatory sequences serving host cell functions (*1*, *8*).

While the cooption of TE sequences is a widespread and recurrent phenomenon that has fueled biological innovation over macroevolutionary timescales (*9*, *10*), very little is known about the early stages in the process. In most cases, the coopted TEs have been traced to ancient TE families that have long been transpositionally inactive. Can active TE families serve essential functions? A small but growing number of studies have identified young, active TEs playing apparently important cellular roles, raising the possibility that TEs and their hosts can engage in mutually beneficial or cooperative relationships (*11–19*). However, these ideas remain contentious in part because it is technically challenging to rigorously test the functionality of active TEs due to their multi-copy and polymorphic nature (*20*, *21*). More broadly, we lack a general framework for understanding how active TEs become integrated into developmental or physiological pathways, where they may exert beneficial activities. To begin addressing these questions within the scope of vertebrate development, we turned to zebrafish, *Danio rerio,* a tractable model for the study of embryogenesis that is host to a plethora of recently active TEs (*22*).

## Results

### Bik-1 is an endogenous retrovirus likely transpositionally active in zebrafish

Previously, we noted that several zebrafish TE families are expressed in embryonic somatic cell lineages, raising the possibility that some of their activities may contribute to normal development (*22*). Here, we focus on BHIKHARI (Bik-1) and BHIKHARI-2 (Bik-2, also known as *crestin*), two related families of endogenous retroviruses expressed in the mesendoderm and neural crest (NC) lineages, respectively. Historically, both Bik-1 and Bik-2 mRNAs have been used as markers of their respective embryonic cell lineages, but to our knowledge the potential developmental role of their encoded products has not been investigated (*23–26*).

Bik-1 is one of five BHIKHARI families annotated in the zebrafish reference genome, all of which are endogenous relatives of the recently discovered lokiretrovirus clade (*27*) (Bik-1-5, Supp. Fig. 1A). The consensus sequence of Bik-1 consists of an internal region flanked by long terminal repeats (LTRs). It contains a single long open reading frame (ORF) predicted to encode a 617-amino acid protein whose origin and putative function have remained uncharacterized (*23*). Our structural prediction analysis predicts that it encodes a Gag protein that likely forms the retroviral capsid (Fig. 1A; Supp. Fig. 2). Bik-1 Gag contains an amino-terminal coiled-coil (CC) domain resembling that of foamy viruses (*28*), a capsid assembly (CA) domain that is structurally conserved throughout retroviral Gag proteins, and a putative carboxy-terminal glycine-rich patch that may be involved in nucleic acid binding (*29*) (Fig. 1A).

**Figure 1.**
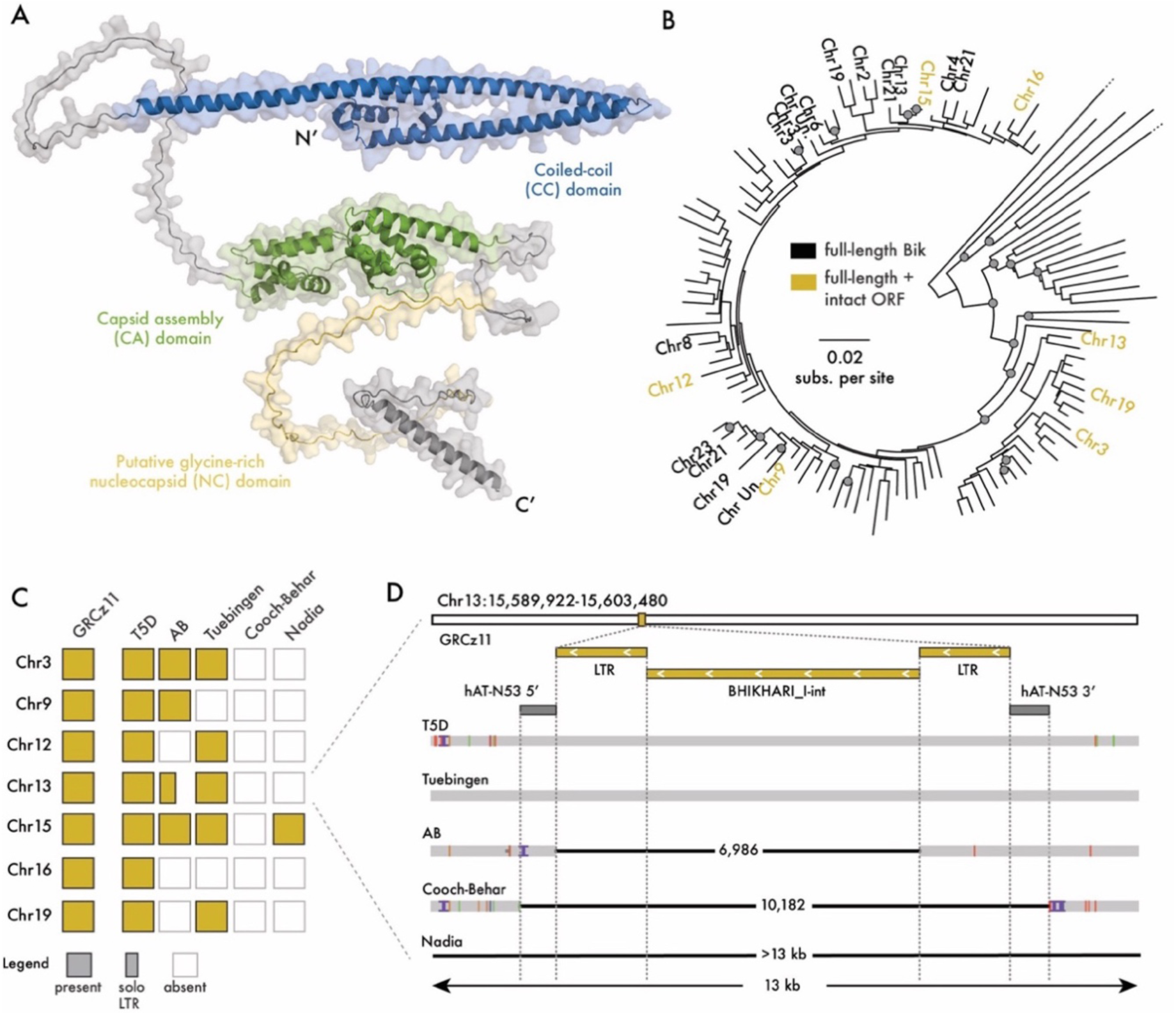
BHIKHARI is a recently active endogenous lokiretrovirus. A) Predicted structure of Bik-1 Gag (via AlphaFold 2.0. (30)) showing conserved structural domains. B) Maximum likelihood phylogenetic tree of de-fragmented Bik-1 insertions between 1000bp and 8750bp in length. Full-length insertions (n= 22, labelled) are defined as those containing internal sequence and both flanking LTRs, and these are further sub-divided into those containing fully intact Gag ORFs (n=7, yellow labels). Gray node labels indicated Shimodaira– Hasegawa approximate likelihood ratio test support ≥ 85 and ultra-fast bootstrap support ≥ 95 (31). C) Analysis of presence/absence insertion polymorphisms from five Danio rerio genome assemblies for the seven fully intact Bik-1 copies in the reference (GRCz11) genome. D) Detailed view of insertion polymorphisms at a Bik-1 locus on Chr13. The reference genome, T5D and Tübingen strains share a full-length insertion, which has been converted to a solo LTR in AB strain. It is located within a previously deposited hAT-N53 insertion, which is itself polymorphic between Cooch-Behar and aforementioned strains. In Nadia, the transpositional history cannot be determined due to a larger deletion of the entire locus.

To assess the transpositional history of Bik-1 in *Danio rerio*, we carried out a comprehensive analysis of Bik-1 insertions in the reference genome (GRCz11). Using TE annotations from a previous analysis (*22*), we found a total of 196 Bik-1 insertions, including 25 “proviral” copies containing both LTRs, and 119 “solo LTRs” formed by ectopic recombination between proviral LTRs. For downstream analysis, we removed fragmented insertions less than 1,000 bp in length, or those with large internal rearrangements. We aligned the resulting 121 DNA sequences and generated a maximum likelihood phylogenetic tree (Fig. 1B, Supp. Data 1). This analysis revealed that many Bik-1 copies are nearly identical across their entire length to their closest neighbor (branch lengths < 0.01 substitutions per site), implying that Bik-1 has replicated very recently. Amongst these are eight proviral copies predicted to encode a full-length Gag protein, which are on average 94% identical to one another at the amino acid level. None of the Bik-1 copies in the reference genome are predicted to encode components of a *pol* gene, such as reverse transcriptase or integrase, implying that transposition of Bik-1 elements must have been complemented by a partner element encoding these activities, possibly an autonomous Bik-1 copy segregating at low frequency in *Danio rerio* populations.

To confirm that Bik-1 has recently transposed, we next looked for polymorphic insertions across *Danio rerio* strains. We generated pairwise genome alignments between the zebrafish reference genome (Tübingen strain) and five additional genome assemblies from lab-raised and wild *Danio rerio* strains (AB, T5D and a second Tübingen for the former, Cooch-Behar and Nadia for the latter). Of the eight proviral insertions with intact Gag ORFs in the reference genome, all were polymorphic between strains (Fig. 1C-D; Supp. Data 2), including precise insertion polymorphisms with perfect target site duplications at the insertion site, a hallmark of recent transposition events (Supp. Fig. 1B-D). Taken together, these data indicate that Bik-1 has mobilized very recently in the germline of *D. rerio* and may still be transpositionally active.

### Efficient depletion of Bik-1 Gag protein using locked nucleic acid antisense oligonucleotides

Bik-1 mRNA is most abundant in adaxial cells, anterior notochord, and prechordal plate of the zebrafish embryo, a pattern driven by transcription of at least half of Bik-1 genomic loci (*22*). To test whether Bik-1 expression is required for zebrafish development, we used a locked nucleic acid (LNA) antisense oligonucleotide to deplete Bik-1 mRNAs. The Bik-1 LNA was designed to target the LTRs of Bik-1, immediately downstream of the previously identified transcription start site (*23*) (Fig. 2A). Allowing for zero sequence mismatches, we predict that the LNA targets at least 114 out of 196 copies of Bik-1 in the reference genome, including 19 of the 25 proviral copies and all of those with a full-length Gag ORF.

**Figure 2.**
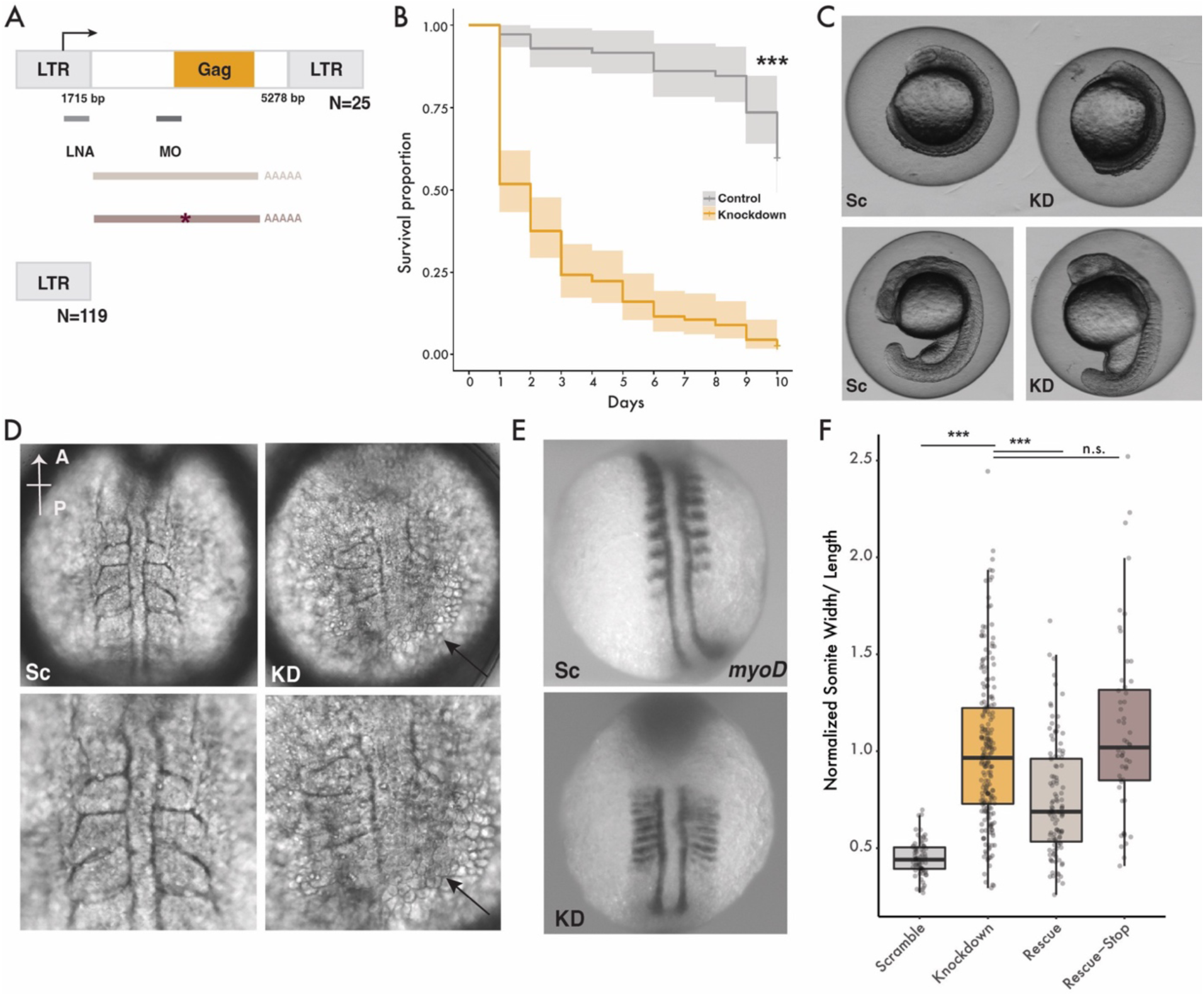
Bik-1 is required for zebrafish somitogenesis. A) Sequence feature of Bik-1 and the experimental design. LNA and MO position indicates their target sequences, respectively. The synthesized Bik-1 mRNA and the one with one nucleotide mutation. B) Kaplan-Meier curve for embryos injected with either Bik-1 LNA or scramble-LNA for 10 continuous days after injection, p-values < 0.0001. C) Phenotypes of embryos injected with Bik-1 LNA or scramble-LNA at 12 (top) or 16 (bottom) hpf. Sc: scramble-LNA, KD: Bik-1 LNA. D) Cell dissociation phenotype (indicated by arrows) in Bik-1 LNA knockdown embryos. E) myoD staining from in situ hybridization at 12 hpf embryos. F) Normalized ratios of somite width/length across multiple injections and treatments, as exemplified by panel E (Mann-Whitney U test, p-value < 0.0001)

After injecting Bik-1 LNA or a scrambled-LNA control into fertilized zygotes, we performed RNA-seq and found that Bik-1 LNA injection resulted in at least 70% reduction in Bik-1 mRNA transcript levels in embryos at 8, 11, 14 and 18 hours post fertilization (hpf) – time points overlapping the endogenous expression of Bik-1. While Bik-1 LTR mRNA was the most significantly down-regulated transcript in Bik-1 LNA injected embryos, 104 host gene transcripts were also differentially expressed (29 up- and 75 down-regulated) by at least two-fold upon Bik-1 knockdown (Supp. Fig. 3, Supp. Data 3). Of these, two protein coding genes (*adka* and *gid8b*) and one non-coding RNA (*CR293502.1*) correspond to downregulated genes located within 10 kb of Bik-1 insertions, suggesting possible cis-regulatory effects associated with Bik-1 knockdown. GO-term enrichment analyses of differentially expressed genes did not reveal significant enrichment for any biological processes or functions. Together these data suggest that Bik-1 LNA efficiently depletes Bik mRNA with limited effects on the rest of the embryonic transcriptome.

### Bik-1 Gag protein is required for zebrafish mesoderm development

Next, we examined the developmental phenotypes of Bik-1 LNA-injected embryos compared to scrambled-LNA controls. We tested a range of LNA concentrations from 8 to 194 pg and found 11 pg of Bik-1 LNA to be the highest concentration that we could inject without immediate lethality. In a survival assay, we found that over 90% of embryos injected with 11 pg Bik-1 LNA passed the shield stage but died within 10 days post-injection, compared to 40% of those injected with the scrambled-LNA control (Fig. 2B), suggesting that Bik-1 expression is essential for zebrafish embryogenesis. Early in development (12 hpf), we observed striking developmental defects for at least 80% of Bik-1 LNA injected embryos, including shortened anterior-posterior axis, microcephaly, laterally extended somites, and cell dissociation (Fig. 2C-D).

To better quantify the somite phenotype, we performed *in situ* RNA hybridization for *myoD*, a somite marker (*32*). Using this labeling, we measured the length normalized to the width of the 2^nd^-5^th^ left-side somites of each embryo. The somites of Bik-1 LNA embryos were significantly longer and narrower than control embryos, indicating a failure of the somite convergence and extension process (*33–35*) (Fig. 2E-F). Throughout convergence and extension, adaxial cells (slow muscle precursors) undergo characteristic morphological changes (*36*, *37*). Bik-1 expression in adaxial cells is amongst the highest of any cell type in the developing embryo (*22*), and following Bik-1 LNA injection we observed clear changes in their organization and morphology (Supp. Fig. 4), further supporting the notion that depleting Bik-1 mRNA impairs somitogenesis.

To corroborate the findings from Bik-1 LNA injection, we designed a morpholino (MO) antisense oligonucleotide to block Bik-1 mRNA translation. The MO sequence was predicted to target at least 17 Bik-1 copies in the reference genome, including all proviral copies with full-length Gag ORF. Bik-1 MO injection recapitulated the developmental defects seen with Bik-1 LNA knockdown, including longer somites and narrower inter-somite spacing (Supp. Fig. 5). Notably, this phenotype strongly resembles that seen in *cdh2* (N-cadherin) and *vangl*2 mutants, in which loss of cell-cell adhesion and cell polarity causes a similar failure of the convergence and extension process (*38*, *39*).

To validate the Bik-1 knockdown phenotype, we attempted to rescue somitogenesis by co-injecting an *in-vitro* transcribed mRNA containing the Bik-1 *gag* ORF preceded by most of its 5’ UTR but excluding the LNA binding site (Fig. 2A). Co-injecting this synthetic Bik-1 mRNA (77 pg) with the Bik-1 LNA (11 pg) partially rescued the somite phenotype (Fig. 2F), indicating that somitogenesis defects in Bik-1 knockdown embryos are caused by the depletion of functional Bik-mRNA. To ascertain whether the phenotype was caused by depletion of the mRNA or its protein product, we co-injected a mutated version of the Bik-1 rescue mRNA with a single nucleotide change introducing a premature stop codon at Leu-69 of the encoded Gag protein. This mutated mRNA failed to rescue the phenotype (Fig. 2F), indicating that the defects in somitogenesis are caused by depletion of Bik-1 Gag protein. Taken together, these experiments indicate that the Gag protein encoded by Bik-1 – a recently active endogenous retrovirus family – is required for early embryonic development in zebrafish.

### Bik-1 Gag associates with the cell membrane

Given the ability of retroviral Gag proteins to target membranes, along with the similarity between the Bik-1 knockdown phenotype and that of mutants for N-cadherin, a membrane-associated protein, we hypothesized that Bik-1 Gag exerts its developmental effects through interaction with the cell membrane. We therefore investigated the subcellular localization of Bik-1 Gag. We overexpressed an HA-tagged Bik-1 Gag in human embryonic kidney 293 (HEK293) cells (Fig. 3A). Through cell fractionation, we found Bik-1 Gag-HA is enriched within the plasma membrane compartment in HEK293 cells relative to the cytosolic fraction, similarly to N-cadherin (Fig. 3B-C). Deletion of the N-terminal CC domain of Bik-1 Gag-HA resulted in the loss of membrane association, and a deletion of the CA domain reduced it (Fig. 3B-C). These results indicate that Bik-1 Gag targets the plasma membrane through its CC domain, and that the CA domain contributes to this activity, possibly through facilitating oligomerization at the cell membrane, as is the case for infectious retroviruses such as HIV (*40*).

**Figure 3.**
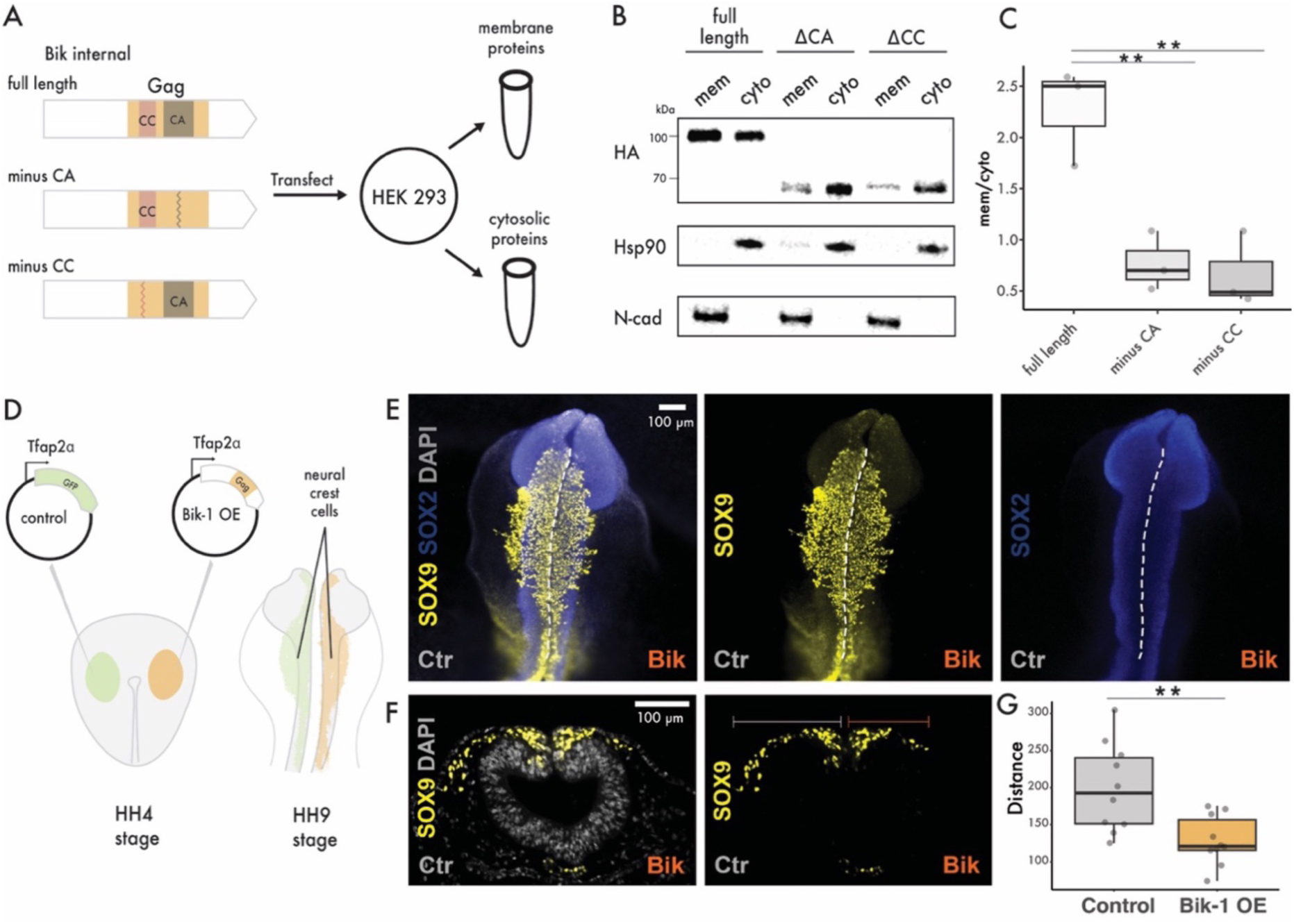
Bik-1 Gag is membrane bound and affects cellular behavior in chicken embryos. A) Experimental design for detecting sublocalization of Bik-1 Gag in HEK293 cells. B) Western blot shows Bik-1 Gag, Bik-1 Gag without CC domain and Bik-1 Gag without CA domain protein in membrane and cytosolic fractions from HEK293 cells. C) Quantification of the blots shown in B (t-test, p=0.0096 and 0.0098 for full length and minus CA or minus CC comparison, respectively). D) Experimental design for overexpression of Bik-1 Gag in chicken embryos. Either GFP or Bik-1 Gag under Tfap2α promoter was injected at HH4-5 stage. We then collected embryos at HH9 stage for downstream analysis. E) Immuofluorescence staining on whole chicken HH9 embryos. SOX9 stains for NC and SOX2 stains for neural tubes. F) Immuofluorescence staining on transverse sections of chicken embryos as shown in D. G) Quantification of NC cell migration distance. Each point represents the average of 3 sections from one embryo. (Wilcoxon rank sum test, p=0.005).

### Bik-1 Gag affects cell migration in chicken embryos

Considering the observed membrane association of Bik-1 Gag in human cells and the phenotypic similarity between Bik-1 knockdown and genetic mutants affecting cell adhesion/interaction, which are evolutionarily conserved processes, we hypothesized that Bik-1 exerts its function through intrinsic, non-specific properties of its Gag protein. We therefore reasoned that ectopic expression of Bik-1 Gag in a heterologous embryo might affect cell migration and development. To test this, we introduced Bik-1 Gag in the developing NC of chicken embryos, a well-characterized experimental paradigm for developmental cell migration. Furthermore, the chicken genome does not appear to host endogenous lokiretroviruses or other elements closely related to Bik-1, enabling us to study the effect of introducing Bik-1 Gag in a naïve host. Overexpressing Bik-1 Gag in chicken embryos at Hamburger and Hamilton (HH) stage 4-5 under the control of a ubiquitous promoter (*β-actin*) caused consistent defects in neural tube closure compared to control embryos (observed in 6/6 embryos). These defects were slightly, but not significantly milder (10/14 embryos) when Bik-1 Gag was expressed using a NC-specific promoter (*Tfap2α*, Supp. Fig. 6). When Bik-1 Gag was only expressed in NC cells, it reduced NC cell migration of those cells (Fig. 3D-G), as is seen in chicken embryos overexpressing N-cadherin (*41*). Thus, whether directly through NC cell migration or indirectly through defects in neural tube closure, Bik-1 Gag affects cell migration in a cell-autonomous fashion and in a heterologous species and tissue.

The effects of Bik-1 Gag overexpression in the developing chicken NC are reminiscent of the activities of another TE-encoded protein called ERNI, which is endogenously expressed in the chicken NC and neural plate (*42–44*). Specifically, overexpressing ERNI has been shown to reduce NC cell migration (*42*). While ERNI’s function and origin have remained elusive, it was previously shown to be encoded by an endogenous retrovirus called ENS-3 (aka Soprano) or its nonautonomous variant, ENS-1, both of which are dispersed throughout the chicken genome (*45*, *46*). Our phylogenetic analysis of the reverse transcriptase domain of ENS-3 confirmed that it is an endogenous retrovirus that forms a sister clade to the ERV-L clade, which includes the previously studied human HERVL and murine MERVL retroelements (*47*) (Supp. Fig. 7A, Supp. Data 4). Structural predictions reveal that ERNI adopts the typical fold of a retroviral capsid domain preceded by an extended CC domain at the N-terminus and a disordered C-terminal region (Supp. Fig. 7B). Overall, the predicted structure of ERNI and Bik-1 Gag are similar, despite being encoded by distantly related endogenous retroviruses. Through phylogenetic analyses, we found that ENS-1/3 is also a very young family, with many full-length, near-identical copies in the chicken genome (median branch length of jungle fowl ENS-1 proviral copies = 0.006 substitutions per site; Supp. Data 4). The number of full-length ENS-1 insertions is also highly variable across chicken breeds, ranging from 4 in Yeonsan Ogye to 44 in Junglefowl, indicating that ENS-1 may have transposed since the domestication of chicken, ∼3,500 years ago (*48*). Thus, like Bik-1, ERNI is a Gag protein encoded by a recently active endogenous retrovirus family with a developmental role in embryonic development.

### Bik-2 Gag protein is required for neural crest cell migration in zebrafish

Having established that Bik-1 is essential for mesoderm development, we turned to Bik-2, which is the closest relative of Bik-1 in zebrafish, but is expressed in the developing NC, rather than the mesendoderm. Like Bik-1, Bik-2 is a nonautonomous family that encodes a *gag* gene but neither *pol* nor *env*; while the amino acid identity between the pairwise alignment of Bik-1 and Bik-2 Gag consensus sequences is only 60%, their predicted structures are nearly identical (Fig. 4A). We identified 393 Bik-2 insertions in the reference genome, of which 337 are solo LTRs. Of the remaining insertions, only five are predicted to encode a full-length Gag protein (Fig. 4B, Supp. Data 1); these insertions are highly similar in sequence to one another, with a mean of 97% amino acid identity between their predicted Gag proteins. Like Bik-1, these intact Bik-2 insertions are insertionally polymorphic between *Danio rerio* strains (Supp. Fig. 8, Supp. Data 2), indicating that Bik-2 has transposed recently.

**Figure 4.**
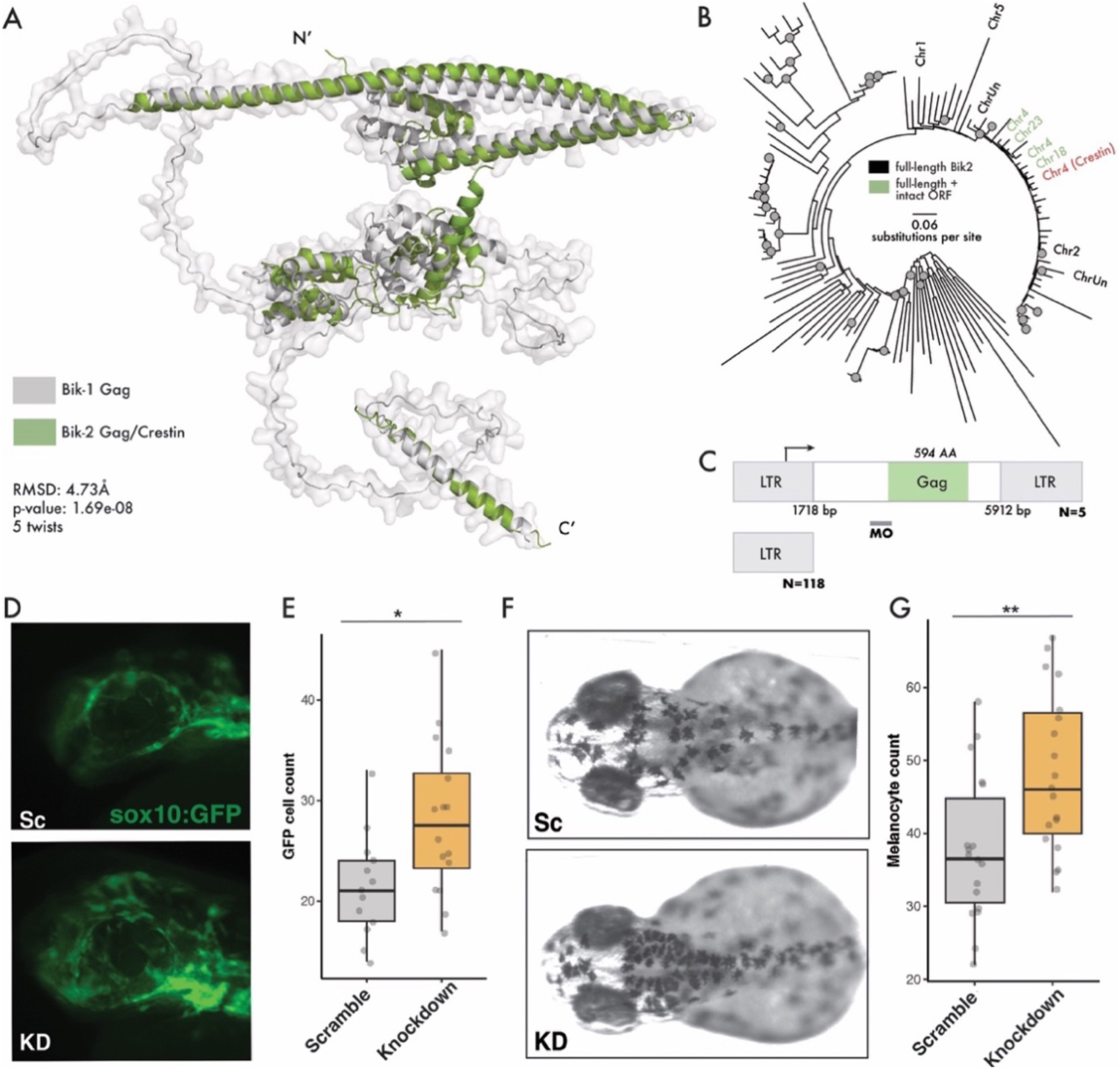
Bik-2 is required for proper migration of neural crest cells. A) Structural alignment of Bik-1 Gag and Bik-2 Gag using FatCat 2.0 (49). B) The Danio rerio reference genome contains 11 near full-length Bik-2 copies (labeled) 4 of which contain an intact Gag open reading frame. Gray nodes indicate ultra-fast bootstrap support > 95 and Shimodaira– Hasegawa approximate likelihood ratio test support > 0.85.; root mean square deviation 4.73Å. C) Sequence features of Bik-2 and the experimental design for MO knockdown. D) sox10:EGFP labeled cells in the head area. E) Numbers of GFP labeled cells in the eye area (Mann-Whitney U test, p-value=0.01388). F) Images of head region of embryos at 54 hpf. G) Melanocyte counts at head region for Bik-1 LNA and scramble-LNA embryos at 54 hpf (Mann-Whitney U test, p-value=0.005912).

In the zebrafish reference genome, the *Gag* gene of one of the Bik-2 elements on Chromosome 4 is annotated as the *crestin* gene, a widely used marker of NC cells (*24–26*). Despite the long-standing use of *crestin* mRNA as a NC marker, its role in development has not yet been characterized. Mining publicly available single-cell RNA-seq data (*22*, *50*), we found that the *crestin* expression pattern is indistinguishable from the rest of the Bik-2 copies in the genome, indicating that *crestin* is merely one of several Bik-2 *gag* genes in the zebrafish genome. Furthermore, the Bik-2 copy carrying the *crestin* gene is also insertionally polymorphic across *D. rerio* strains (Supp. Fig. 8).

To determine whether Bik-2 Gag protein plays a role in NC development, we designed a MO to block Bik-2 Gag translation in a transgenic *sox10:EGFP* zebrafish line enabling visualization of NC cells (*51*) (Fig. 4C). We observed abnormal migration patterns of NC cells in the Bik-2 MO injected embryos compared to scrambled MO embryos. To quantify this phenotype, we focused on the eye area of zebrafish at 24 hpf because it normally contains very few NC cells at this stage and has a definable boundary (*52*). Counting the number of GFP-labeled cells, we found that Bik-2 MO-injected embryos have significantly more NC cells in the eye area compared to controls (Fig. 4D-E). In later developmental stages, NC cells differentiate into melanocytes; thus, if Bik-2 MO perturbs NC migration it may also affect the position and number of melanocytes through the body. Indeed, we found that Bik-2 MO-injected embryos at 54 hpf have significantly more melanocytes in the head area compared to controls (Fig. 4F-G). Together, these data indicate that Bik-2 Gag protein is required for proper NC development in zebrafish.

## Discussion

In this work, we have demonstrated that the expression of Gag proteins encoded by TEs recently active in zebrafish is required for proper embryonic development. Bik-1, an endogenous lokiretrovirus with dozens of near-identical, polymorphic insertions in the zebrafish genome, encodes a Gag protein essential for somitogenesis. Bik-2, a distant relative of Bik-1, encodes a structurally similar Gag protein, but is specifically expressed in zebrafish NC cells. Depleting Bik-2 Gag perturbs NC development, indicating that expression of Bik-2 Gag is also important for embryogenesis, despite operating in a different cell lineage than Bik-1.

Several lines of evidence suggest that Bik-1 and Bik-2 modulate cell adhesion and/or migration. First, Bik-1 accumulates at the plasma cell membrane when overexpressed in human cells. Second, the phenotype of Bik-1 knockdown resembles phenotypes of genetic mutants for cell adhesion proteins, such as N-Cadherin. Third, both depletion of Bik-2 in zebrafish and ectopic expression of Bik-1 Gag in chicken causes aberrant NC cell migration. The fact that Bik-1 Gag protein exerts this activity when expressed ectopically in a heterologous species and cell type suggests that its developmental effects are mediated by intrinsic features of the Gag protein.

These effects may not be restricted to the zebrafish BHIKHARI family. Indeed, ERNI, a previously studied NC regulator in chicken, is also a Gag protein encoded by an active endogenous retrovirus, ENS-1. Thus, three different vertebrate endogenous retroviruses encode Gag proteins that are critical for normal development. Notably, ENS-1 is related to MERVL, another active element which may be required for mouse preimplantation development, albeit through an as yet poorly understood mechanism (*14*, *17*, *53*). Moreover, many other Gag proteins from diverse LTR retroelements have repeatedly been coopted throughout eukaryotic evolution, including examples such as Arc and Peg10 (*54–56*).

Our findings add to recent groundbreaking studies suggesting that some active TEs are not strictly selfish, but instead cooperate with their host to fulfill beneficial or even essential biological processes such as embryonic development (*11–19*). This is unexpected because developmental processes like somitogenesis or NC development rely on deeply conserved pathways across vertebrates, while active TEs such as Bik-1, Bik-2 and ENS-1 have recently spread within populations. We propose an “addiction” model to reconcile this paradox and explain how the products of young TEs can be incorporated into conserved cellular pathways, potentially paving the way toward their complete domestication (Fig. 5).

**Figure 5.**
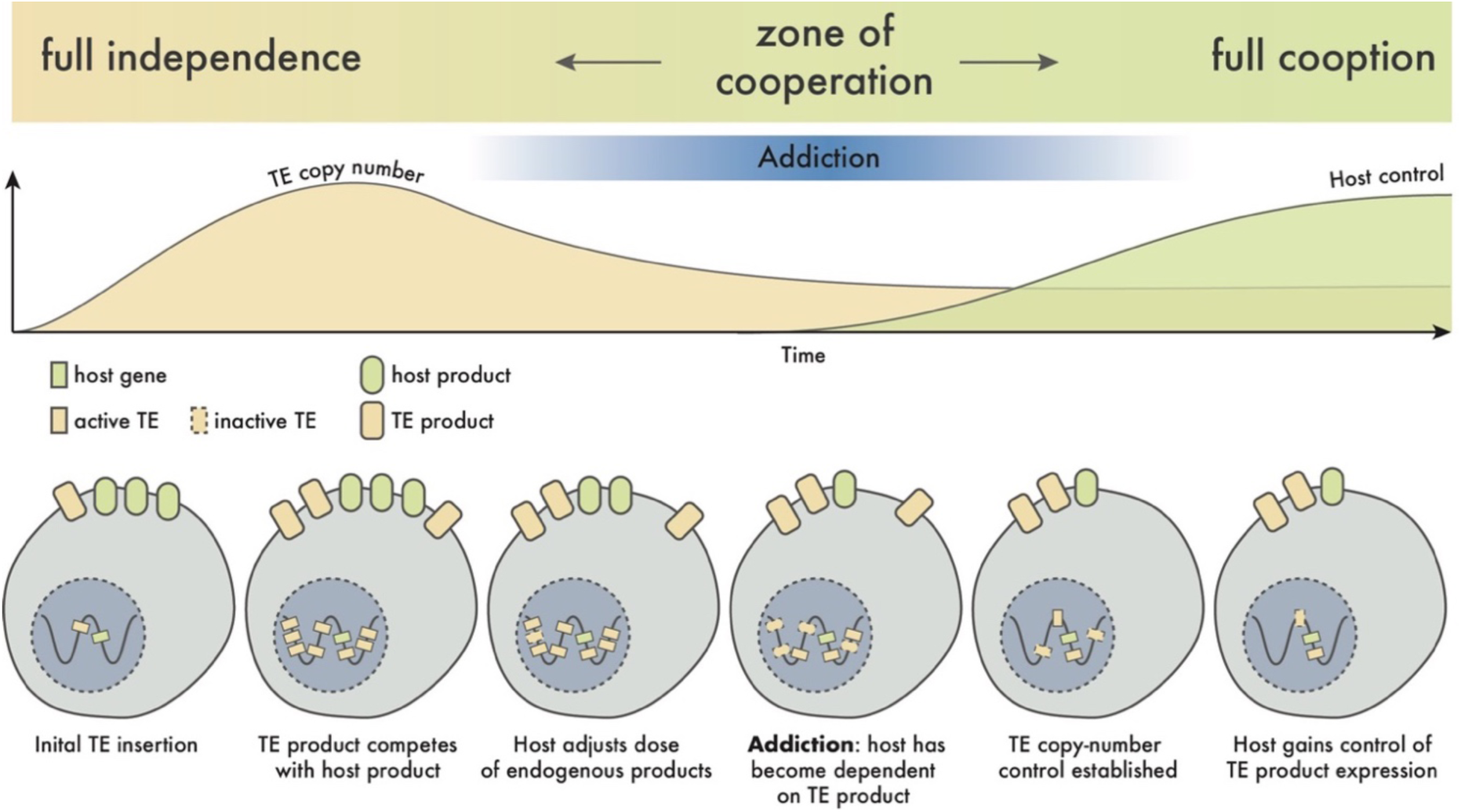
A model of host-TE cooperation through addiction. As TE copy number increases, interaction between host and TE products, whether antagonistic or beneficial (e.g. TE product serves redundant functions), may lead to the host becoming addicted. This situation would force cooperation between host and TE, until such time as the TE becomes inactive or the host gains control over copy number.

We envision that certain TEs encode products with cellular activities that encroach on or are functionally redundant with host products, e.g. Gag redundant with cell adhesion molecules. Much like gene duplicates, the functional redundancy introduced by every new TE copy expressing the product may relax the reliance and evolutionary constraint on the host-encoded product. As the genomic TE copy number increases, so too will host dependency on the TE product. Eventually, an unstable equilibrium is reached when the host becomes effectively addicted to the TE product. This process may benefit both host and TE by establishing a form of TE copy number control (*5*), as a sudden increase in the dose of TE product would likely be toxic to the host. As TE copy number stabilizes, the host may kick the addiction by taking control of the supply of the product, either through a trans-regulatory mechanism such as piRNAs, as observed for ERNI/ENS-1 in chicken (*42*), or by taking cis-regulatory control over a specific TE locus, thereby paving the way toward a coopted TE gene.

Our understanding of the nature of TEs and their interactions with their hosts has undergone multiple paradigm shifts over the last half-century, from their first visionary description as “controlling elements” (*57*), through the “junk DNA” era that followed the sequencing of the first complete genomes (*58*). Over the past half-century, the prevailing view of TEs has been that they are deleterious parasites that, once immobilized, by virtue of their diversity and abundance, provide a source of raw genetic material which may be repurposed for organismal function. However, the findings presented here and elsewhere (*11–19*) are challenging this view by showing that young, active TEs can establish cooperative interactions that make them indispensable to their host organism.

## Materials and Methods

### Zebrafish husbandry

Zebrafish were maintained in the facility of Prof. Joseph Fetcho’s lab at Cornell University. All animal work was performed under IACUC protocols and in accordance with the U.S. National Institutes of Health guidelines for animal use in experiments. All experimental procedures involving zebrafish were approved by Cornell University’s Institutional Animal Care and Use Committee.

### Phylogenetic analysis of BHIKHARI family members

To investigate the evolutionary origin of the BHIKHARI superfamily, we identified related endogenous lokiretroviral sequences in the RepBase v28.08 library of consensus TE sequences (*59*) using blastn with the consensus sequences for internal regions of Bik-1-5 (*60*). Using the same process, we identified close relatives of Bik-1-5 in other Danio species by searching de-novo TE libraries generated using RepeatModeler2 and the -LTRStruct module (*61*). The resulting set of sequences was supplemented with the internal sequences of foamy retroviruses from the NCBI virus database and Odinretroviruses from a recently published paper describing the family (*62*).

Gag and Pol amino acid sequences were extracted and aligned separately using MAFFT in einsi mode (gap extension penalty = 0) (*63*). After concatenating the Gag and Pol alignments, a partitioned phylogenetic tree was calculated from this alignment using IQ-Tree v2.2.0 with ModelFinder (*64*, *65*). Ultra-fast bootstrap and SH-aLRT support values were generated with 1000 bootstrap replicates (*31*). The resulting tree was rooted using the foamy retroviruses as an outgroup.

To assess the age of Bik insertions in the Danio rerio reference genome, we re-used previously reconstructed (LTR+internal+LTR) insertion sequences (*22*). To avoid dealing with highly-fragmented sequences, we filtered this set to those between 1000 and 8750bp in length, thus allowing for partially-intact solo LTRs, but removing full-length elements with large insertions or internal rearrangements. We aligned these sequences using MAFFT, then manually edited the resulting alignment to remove spuriously aligned regions. A terrace-aware partitioned phylogenetic analysis was used to generate a tree from this alignment, with LTRs and internal sequences modelled separately (*66*). We carried out an identical process to generate the Bik-2 tree, with the sole difference being that we restricted sequences to those between 1000 and 9350bp in length, reflecting the larger size of Bik-2 elements.

### Analysis of insertion polymorphisms

Presence/absence insertion polymorphisms for transpositionally competent copies of Bik-1/2 in the reference genome (GCF_000002035.6) were identified in five additional *Danio rerio* genome assemblies (GCA_903684865.1: AB; GCA_903684855.2: Tuebingen; GCA_018400075.1: T5D; GCA_903798165.1: Cooch-Behar; GCA_903798175.1 – Nadia). For each of these five genomes, minimap v2.24 with the -asm5 flag was used to align the target assembly to the reference (*67*). a region of the genome containing the locus and 50kb of flanking sequence was extracted using BEDtools v2.30.0 and SAMtools v1.18 (*68*, *69*). The resulting BAM files were manually inspected to determine the polymorphism status at each Bik-1/2 locus.

### Structural analysis of Gag proteins

To predict the monomeric structure of Bik-1 Gag, we used AlphaFold2 via the ColabFold notebook (*30*, *70*). We compared the CA domain of Bik-1 to that of three other Gag proteins (PDB: 6TAP-D, 6OMT-A, 5A9E-A; (*71–73*)). To flexibly align these structures, we used FATCAT 2.0 to allow for twists in the structure of linker sequence (*49*). For Bik-2 Gag, we used the pre-computed structure of Crestin available from the AlphaFold protein structure database (*74*), and aligned this to Bik-1 using FATCAT 2.0. Finally, the predicted 3D structure of ERNI was obtained from the AlphaFold database and the CA domain was aligned to that of a foamy retrovirus Gag protein (PDB: 51MHl; (*75*)) using CEalign (*76*) in PyMOL (*77*).

### Knockdown of Bik-1

To target Bik-1 transcription, 22nt locked nucleic acids (LNAs) targeting the transcription start site of Bik-1 and a scrambled control were synthesized by IDT (5’GGTGACGACTGGCAAAGATGAC and 5’GGGATATTACAAGCGGACGCAG, respectively). 22 pg of either Bik-LNA or scramble-LNA were injected into zebrafish embryos in fertilized zygotes at the one-cell stage. Embryos were collected at 8, 12, 16 and 18 hpf for downstream experiments. To target translation, a 25nt MO antisense oligonucleotide (GTCTTTGGGGTGAGCCATGTTTTGC) and a scrambled control were synthesized by Gene Tools, LLC. 2.7ng of MO was injected into zebrafish embryos at one-cell stage. Embryos were collected at 12 hpf for downstream analysis.

### Total RNA sequencing and analysis

9-11 embryos injected with either Bik-1 LNA or scrambled LNA were collected at 8, 12, 16 and 18 hpf. Two biological replicates for each time point per treatment were performed. Total RNAs were extracted using Qiagen RNeasy kit and sent to Novogene for stranded RNA sequencing. Raw reads were filtered using Trimmomatic and aligned to the reference genome using STAR v2.11a (*78*, *79*). Differential expression analysis was performed using DE-seq2 (*80*).

### Bik-1 rescue injections

Bik-1 mRNA containing its 5’UTR and open reading frame (but lacking the Bik-1 LNA target site) was amplified from cDNAs via PCR and cloned into pT3TS-nCas9n vector (addgene #46757, a gift from Ariel Bazzini). Bik-1 mRNA was then synthesized using mMESSAGE mMACHINE™ T3 Transcription Kit (AM1348). The mRNA was purified using RNeasy kit from Qiagen. The Bik-1 mRNA (153 pg) combined with Bik-1 LNA (22 pg) or Bik-1 LNA alone were co-injected into embryos at one-cell stage as the rescue experiment. Injected embryos then collected at 12 hpf for in situ hybridization.

We used In-Fusion enzyme (Takara Bio, #638947) to create a single nucleotide substitution on the 69th amino acid (Leucine to Stop) of Gag. The Bik-1 mRNA with the mutation (Rescue-Stop) was synthesized and purified as above. The Bik-1 mRNA with the mutation (153 pg) combined with Bik-1 LNA (22 pg) or Bik-1 LNA alone were co-injected into embryos at one-cell stage as in the original rescue experiment. Injected embryos were then collected at 12 hpf for in situ hybridization.

### Quantification of the somite phenotype

Injected embryos were collected at 12 hpf and fixed in 4% PFA. In situ hybridization was performed as described (*81*). MyoD probe was amplified by PCR and synthesized as described (*32*). After colorization, pictures of each embryo were taken with a Leica upright microscope. To control for the developmental variation, only embryos with 9-13 somites will be used for downstream quantification. Width and length of 2nd to 5th left somite of each embryo were counted using ImageJ (Fiji) (*82*). A ratio was calculated as width/length for each somite. To account for the variation between injection rounds, each ratio was normalized to the mean ratio of knockdown embryos for each replicate.

### In situ hybridization and immunofluorescence staining in zebrafish embryos

Primers targeting myoD and Bik-1 were used to PCR probes from wild-type cDNAs of zebrafish embryos. Whole-mount in situ hybridization was perform as previous described (*81*). Embryos hybridized with myoD probes were imaged using Leica microscope and measured the somite length/width using ImageJ. Fluorescence in situ hybridization was performed using the same protocol except for the Tyamide Signal Amplification in the end. Embryos were imaged using Zeiss 710 confocal microscope to detect the fluorescence signal.

Embryos injected with Bik-1 LNA or scramble-LNA were collected at 12 or 16 hpf and used for immunofluorescence experiments following the protocol as described (*83*), with primary antibody (F59 from DSHB) and a second antibody (anti-Mouse Alexa Fluor 488 from Thermo Fisher Scientific). Embryos were mounted and imaged using a Zeiss confocal z710 microscope. The shape of adaxial cells and the angle of slow muscle chevrons were quantified using ImageJ.

### Bik-1 expression in HEK293FT cells

Either a full-length, deleted CA (capsid) or deleted CC (coiled-coiled) version of Bik-1 with an HA tag was inserted into the pcDNA3.1(+) backbone (Invitrogen: V79020). HEK293T cells were transfected with Lipofectamine™ 2000 (Invitrogen: 11668019) at a ratio of 5 uL Lipofectamine™ 2000: 2.5 ug plasmid DNA diluted in Opti-MEM (Gibco: 31985062). After 48 hrs, cells were harvested, and the cytosolic and membrane proteins were separated using the Mem-PER Plus Membrane Protein Extraction Kit (Thermo Fisher: 89842) with cOmplete™ Mini Protease Inhibitor Cocktail (Roche: 4693124001). Protein samples were boiled for 10 minutes at 95 °C and loaded onto Mini-PROTEAN TGX™ Precast Gels (BIO-RAD:4569036), and electrophoresis was performed according to manufacturer’s guidelines. Proteins were then transferred to PVDF membranes using iBlot™ 3 Western Blot Transfer Device (Invitrogen: IB31001) with iBlot™ 3 Transfer Stacks (Invitrogen: IB34001, IB34002). After blocking in 5% non-fat milk, blots were incubated with primary antibodies (HA-Cell Signaling: #2367; Hsp90-Thermo Fisher: MA1-10372; N-cadherin-Thermo Fisher: 33-3900) overnight at 4°C. The next day, bots were washed three times using TBST and incubated with secondary antibodies (Cell Signaling: #7074 and #7076). The blots were then visualized and imaged using Pierce ECL Western Blotting Substrate (Thermo Fisher 32106) or SuperSignal™ West Atto Ultimate Sensitivity Substrate (Thermo Scientific: A38554) and a Li-Cor Odyssey imager. Each blot was exposed for three time-points to ensure that the intensity of the bands was in a liner range. Then, the average intensity of each band was measured using ImageJ.

### Bik-1 overexpression in chicken embryos

Fertilized White Leghorn chicken embryos were purchased from the University of Connecticut (Department of Animal Science). The embryos were incubated at 37°C until the desirable stage based on the Hamburger and Hamilton staging system. A full-length zebrafish Bik-1 was inserted into a pTK-EGFP backbone with an avian Tfap2aE1 enhancer (*84*, *85*). 1.25 µg/µl of Tfap2aE1-Bik-1 or an empty vector was injected bilaterally into the space between the epiblast and the vitelline membrane and electroporated with platinum electrodes by passing five 50 msec pulses of 5.2 V. of chick embryos (HH4-5) *ex ovo* (*86*). The embryos were then incubated in a 37 °C humidified chamber until HH10 for the following experiments.

### Immunofluorescence staining in chicken embryos

Transfected embryos were collected at HH10 and fixed in 4% PFA. Subsequently, the embryos were washed in TBST (1X TBS with 0.1% TritonX-100) and TBTD (1X TBST with 1% DMSO). After blocking with 10% Donkey Serum (Equitech-Bio, sd300500) in TBTD for 1 hour, embryos were incubated with primary antibodies (SOX9: Millipore Sigma ab5535 1:200; SOX2: R&D Systems AF2018 1:200) at 4°C overnight. The following day, embryos were washed in TBTD 3 times and incubated with secondary antibodies (Alexa Fluor 488 and 633: Thermo Fisher Scientific, A21206 and A21126, respectively) and DAPI. Whole embryos were imaged using a Zeiss Imager.Z2 fluorescent microscope. For cryosectioning, embryos were transferred to 5% sucrose (Sigma S7903) in 1X PBS for 2 hours at room temperature, followed by 15% sucrose in 1X PBS at 4°C overnight. Embryos were then incubated at 37°C in 7.5% gelatin (Sigma G1890) in 15% sucrose PBS solution for 2 hours and embedded in silicone molds underwent flash frozen in liquid nitrogen. The samples were sectioned using a CryoStar cryotome. The sections were transferred onto charged glass slides at room temperature overnight. The next day, the slides were incubated in 1X PBS at 42°C for 15 min and 1X PBS at room temperature for 10 min 3 times to wash off the gelatin. After mounting with Fluoromount, slides were imaged using a Zeiss Imager.Z2 fluorescent microscope.

### Phylogenetic analysis of ERNI/ENS1

To confirm the phylogenetic placement of ENS1/3, we first assembled a set of retrovirus and LTR retrotransposon Pol amino acid sequences by searching RepBase v28.08 and the NCBI virus database using blast (*60*). Adding ENS3 Pol to this list, we extracted the reverse transcriptase domain using the PFAM HMMs RVT_1 and RVT_2 (PF00078.28 and PF07727.15) and the hmmsearch tool from HMMER v3.3.2 (*87*). The resulting sequences were aligned using MAFFT v7.490 and used to produce a phylogenetic tree via IQ-TREE v2.06, run with 1000 ultra-fast bootstrap and Shimodaira-Hasegawa approximate likelihood ratio test replicates (*31*, *63*, *65*).

To compare ENS1 copy number across chicken strains, we combined the LTR and internal consensus sequences of ENS1 to reconstruct a full-length consensus of ENS1 (4666bp), and used blastn to search for this sequence in ten chicken genome assemblies (GCF_000002315.6, GCF_016699485.2, GCF_016700215.2, GCA_024652985.1, GCA_024652995.1, GCA_024653025.1, GCA_024653035.1, GCA_002798355.1, GCA_024206055.1, GCA_024653045.1), using an e-value threshold of 1e-60, and a minimum and maximum size of 2333 and 5133 to restrict the results to near full-length sequences. To assess the median age of insertions, a phylogenetic tree of near full-length jungle fowl insertions (GCF_000002315.6) was created using MAFFT and IQ-TREE, as described for Bik-1.

### Knockdown of Bik-2

25nt of MO antisense oligonucleotide (GGGAAGACCTATAACTGACACTGAC) and a scrambled control were synthesized by Gene Tools, LLC. 14 ng of MO was injected into zebrafish embryos at one-cell stage. Embryos were collected at 24 hpf for imaging. The GFP-positive cells within the eye were counted using ImageJ.

### Data Access

All raw and processed sequencing data generated in this study will be submitted to the NCBI Gene Expression Omnibus. Scripts used in the generation and analysis of data for this project can be found at GitHub (https://github.com/jonathan-wells/bhikhari).

## Acknowledgments

We thank Dr. Joseph Fetcho for support with zebrafish husbandry and advice throughout the project. We thank the Social Evolution and Behavior course from Rockefeller University for sparking the idea about cooperation. We thank the MBL Zebrafish 2018 course for imparting fundamental technical skills. We thank members of the Feschotte laboratory and Dr. Volker Vogt for valuable feedback and discussion. We thank Sharon Amacher and Chris Fromme for experimental suggestions. This work was supported by grant R35-GM122550 from the National Institutes of Health to C.F. J.N.W. was supported by a Human Frontier Science Program long-term fellowship (LT000017/2019-L). N.-C.C. was supported by a Distinguished Scholar Award from the Cornell Center for Vertebrate Genomics. Author contributions: Initial project conception by N.-C.C. Experimental design by N.-C.C., J.N.W. and C.F. Bik-1/2 perturbation experiments carried out by N.-C.C. Comparative phylogenetic and structural analyses carried out by J.N.W. A.Y.W. performed membrane association assays. P.S. carried out early pilot experiments and phylogenetic analyses demonstrating association of Bik family with lokiretroviruses. N.-C.C, Y.-C.H. and M.S.-C. facilitated experiments in chicken embryos. V.H.T. predicted multimeric Bik-1 Gag structures. N.-C.C, J.N.W. and C.F. wrote the manuscript.

## Notes

https://github.com/jonathan-wells/bhikhari

### Competing Interest Statement

The authors have declared no competing interest.

## References

1. G. Bourque et al., Genome Biol 19, 199 (2018).

2. J. N. Wells, C. Feschotte, Annu Rev Genet 54, 539 (2020).

3. L. M. Payer, K. H. Burns, Nat Rev Genet 20, 760 (2019).

4. J. H. Werren, Proc Natl Acad Sci U S A 108 Suppl 2, 10863 (2011).

5. B. Charlesworth, C. H. Langley, Annu Rev Genet 23, 251 (1989).

6. W. F. Doolittle, C. Sapienza, Nature 284, 601 (1980).

7. L. E. Orgel, F. H. Crick, Nature 284, 604 (1980).

8. E. B. Chuong, N. C. Elde, C. Feschotte, Nat Rev Genet 18, 71 (2017).

9. R. L. Cosby, N. C. Chang, C. Feschotte, Genes Dev 33, 1098 (2019).

10. A. Gebrie, Mob DNA 14, 9 (2023).

11. M. L. Pardue, P. G. DeBaryshe, Annu Rev Genet 37, 485 (2003).

12. J. W. Jachowicz et al., Nat Genet 49, 1502 (2017).

13. M. Percharde et al., Cell 174, 391 (2018).

14. K. Kruse, et al., *bioRxiv* (2019).

15. R. S. Moore et al., Cell 184, 4697 (2021).

16. J. O. Nelson, A. Slicko, Y. M. Yamashita, Proc Natl Acad Sci U S A 120, e2221613120 (2023).

17. A. Sakashita et al., Nat Genet 55, 484 (2023).

18. P. G. M’Angale, et al., *bioRxiv* (2023).

19. S. Cranz-Mileva et al., Genome Biol Evol 16, evae010 (2024).

20. K. O’Neill, D. Brocks, M. G. Hammell, Philos Trans R Soc Lond B Biol Sci 375, 20190345 (2020).

21. F. S. de Souza, L. F. Franchini, M. Rubinstein, Mol Biol Evol 30, 1239 (2013).

22. N. C. Chang, Q. Rovira, J. Wells, C. Feschotte, J. M. Vaquerizas, Genome Res 32, 1408 (2022).

23. A. M. Vogel, T. Gerster, Mech Dev 85, 133 (1999).

24. R. Luo, M. An, B. L. Arduini, P. D. Henion, Dev Dyn 220, 169 (2001).

25. A. L. Rubinstein, D. Lee, R. Luo, P. D. Henion, M. E. Halpern, Genesis 26, 86 (2000).

26. C. K. Kaufman et al., Science 351, aad2197 (2016).

27. J. Wang, G. Z. Han, Mol Biol Evol 38, 1031 (2021).

28. J. Tobaly-Tapiero et al., J Virol 75, 4367 (2001).

29. M. V. Hamann et al., Retrovirology 11, 87 (2014).

30. J. Jumper et al., Nature 596, 583 (2021).

31. D. T. Hoang, O. Chernomor, A. von Haeseler, B. Q. Minh, L. S. Vinh, Mol Biol Evol 35, 518 (2018).

32. E. S. Weinberg et al., Development 122, 271 (1996).

33. N. S. Glickman, C. B. Kimmel, M. A. Jones, R. J. Adams, Development 130, 873 (2003).

34. M. L. K. Williams, L. Solnica-Krezel, Curr Top Dev Biol 136, 377 (2020).

35. A. F. Schier, W. S. Talbot, Annu Rev Genet 39, 561 (2005).

36. D. F. Daggett, C. R. Domingo, P. D. Currie, S. L. Amacher, Dev Biol 309, 169 (2007).

37. C. Yin, L. Solnica-Krezel, Dev Dyn 236, 2742 (2007).

38. M. J. Harrington, E. Hong, O. Fasanmi, R. Brewster, BMC Dev Biol 7, 130 (2007).

39. X. Pan, V. Sittaramane, S. Gurung, A. Chandrasekhar, Mech Dev 131, 1 (2014).

40. Y. Fang et al., PLoS Biol 5, e158 (2007).

41. I. Shoval, A. Ludwig, C. Kalcheim, Development 134, 491 (2007).

42. R. Galton, K. Fejes-Toth, M. E. Bronner, Sci Adv 8, eabn1441 (2022).

43. S. Blanc et al., PLoS One 9, e92039 (2014).

44. C. Papanayotou et al., PLoS Biol 6, e2 (2008).

45. E. Lerat, A. M. Birot, J. Samarut, A. Mey, J Mol Evol 65, 215 (2007).

46. T. Wicker et al., Genome Res 15, 126 (2005).

47. P. G. Hendrickson et al., Nat Genet 49, 925 (2017).

48. J. Peters et al., Proc Natl Acad Sci U S A 119, e2121978119 (2022).

49. Z. Li, L. Jaroszewski, M. Iyer, M. Sedova, A. Godzik, Nucleic Acids Res 48, W60 (2020).

50. J. A. Farrell et al., Science 360, eaar3131 (2018).

51. N. Wada et al., Development 132, 3977 (2005).

52. T. Langenberg, A. Kahana, J. A. Wszalek, M. C. Halloran, Dev Dyn 237, 1645 (2008).

53. S. de la Rosa, M. Del Mar Rigual, P. Vargiu, S. Ortega, N. Djouder, Sci Adv 10, eadk9394 (2024).

54. J. Wang, G. Z. Han, Mol Biol Evol 38, 3267 (2021).

55. E. D. Pastuzyn et al., Cell 172, 275 (2018).

56. M. Segel et al., Science 373, 882 (2021).

57. B. McClintock, The American Naturalist 95, 265 (1961).

58. A. F. Palazzo, T. R. Gregory, PLoS Genet 10, e1004351 (2014).

59. W. Bao, K. K. Kojima, O. Kohany, Mob DNA 6, 11 (2015).

60. S. F. Altschul, W. Gish, W. Miller, E. W. Myers, D. J. Lipman, J Mol Biol 215, 403 (1990).

61. J. M. Flynn et al., Proc Natl Acad Sci U S A 117, 9451 (2020).

62. J. Wang, G. Z. Han, mBio 13, e0018722 (2022).

63. K. Katoh, D. M. Standley, Mol Biol Evol 30, 772 (2013).

64. S. Kalyaanamoorthy, B. Q. Minh, T. K. F. Wong, A. von Haeseler, L. S. Jermiin, Nat Methods 14, 587 (2017).

65. B. Q. Minh et al., Mol Biol Evol 37, 1530 (2020).

66. O. Chernomor, A. von Haeseler, B. Q. Minh, Syst Biol 65, 997 (2016).

67. H. Li, Bioinformatics 37, 4572 (2021).

68. A. R. Quinlan, I. M. Hall, Bioinformatics 26, 841 (2010).

69. H. Li et al., Bioinformatics 25, 2078 (2009).

70. M. Mirdita et al., Nat Methods 19, 679 (2022).

71. P. T. Huang et al., Cell Rep 28, 2373 (2019).

72. F. K. Schur, R. A. Dick, W. J. Hagen, V. M. Vogt, J. A. Briggs, J Virol 89, 10294 (2015).

73. S. Erlendsson et al., Nat Neurosci 23, 172 (2020).

74. M. Varadi et al., Nucleic Acids Res 50, D439 (2022).

75. N. J. Ball et al., PLoS Pathog 12, e1005981 (2016).

76. I. N. Shindyalov, P. E. Bourne, Protein Eng 11, 739 (1998).

77. The PyMOL Molecular Graphics System, Version 3.0 Schrödinger, LLC.

78. A. M. Bolger, M. Lohse, B. Usadel, Bioinformatics 30, 2114 (2014).

79. A. Dobin et al., Bioinformatics 29, 15 (2013).

80. M. I. Love, W. Huber, S. Anders, Genome Biology 15, 550 (2014).

81. C. Thisse, B. Thisse, Nat Protoc 3, 59 (2008).

82. J. Schindelin et al., Nat Methods 9, 676 (2012).

83. D. F. Daggett et al., Curr Biol 14, 1632 (2004).

84. A. S. Hovland et al., Dev Cell 57, 2257 (2022).

85. M. Uchikawa, Y. Ishida, T. Takemoto, Y. Kamachi, H. Kondoh, Dev Cell 4, 509 (2003).

86. M. Simões-Costa, M. Stone, M. E. Bronner, Dev Cell 34, 544 (2015).

87. S. R. Eddy, PLoS Computational Biology 7, e1002195 (2011).

