## Supplementary Figures for "Gag proteins encoded by endogenous retroviruses are required for zebrafish development"

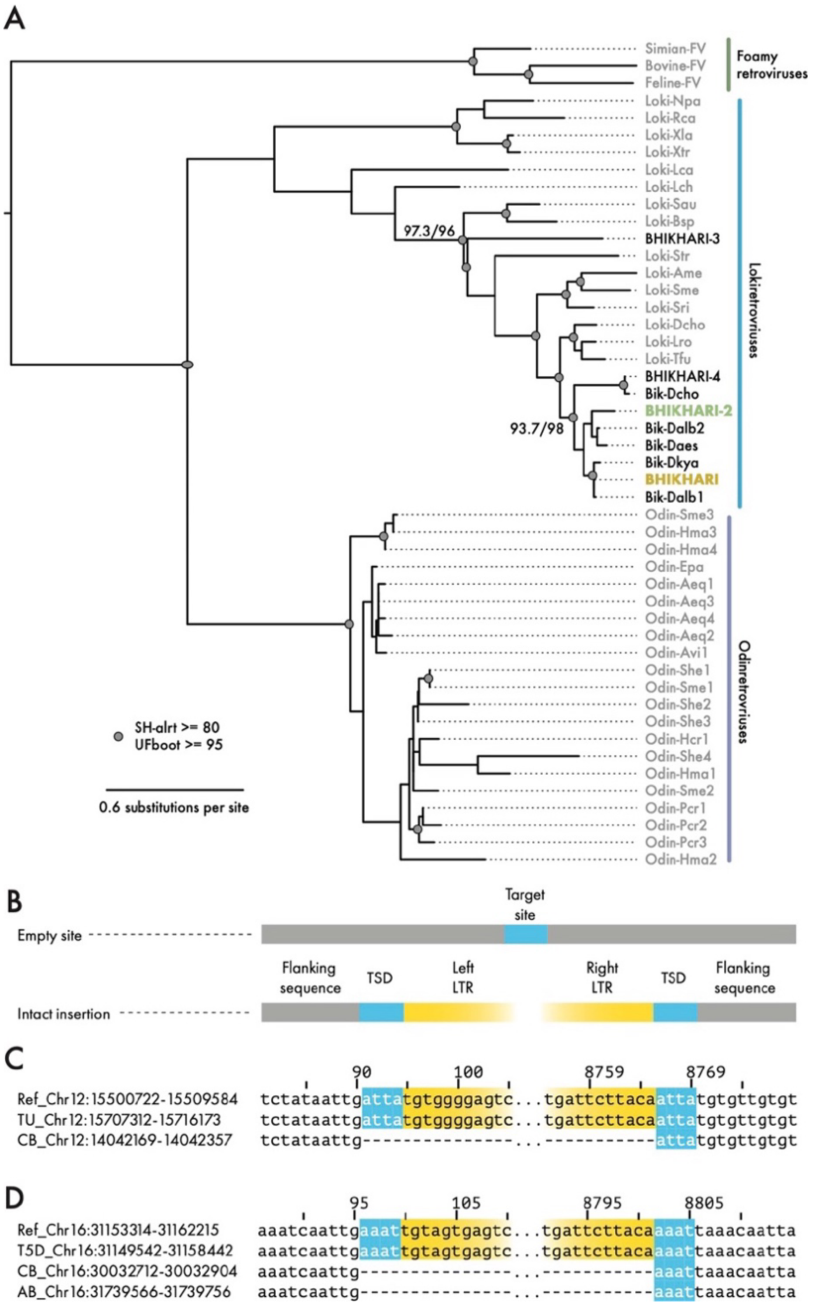


***Supplementary Figure 1. Bik-1 is an endogenous lokiretrovirus with polymorphic insertions***
*A) Maximum-likelihood phylogenetic tree of concatenated Gag CA and Pol reverse transcriptase domains. Reverse transcriptase sequences from foamy retroviruses were used to root the tree. Odinretrovirus sister group sequences accessed from (1). All Danio rerio BHIKHARI family members with detectable coding sequence form a monophyletic clade within lokiretroviridae (Shimodaira–Hasegawa approximate likelihood ratio test support = 97.3; ultra-fast bootstrap support = 96). B) Cartoon showing target site duplications (TSDs) generated by LTR retroelement insertion. C) Full-length polymorphic insertion on Danio rerio Chromosome 12 and D) Chromosome 16.*

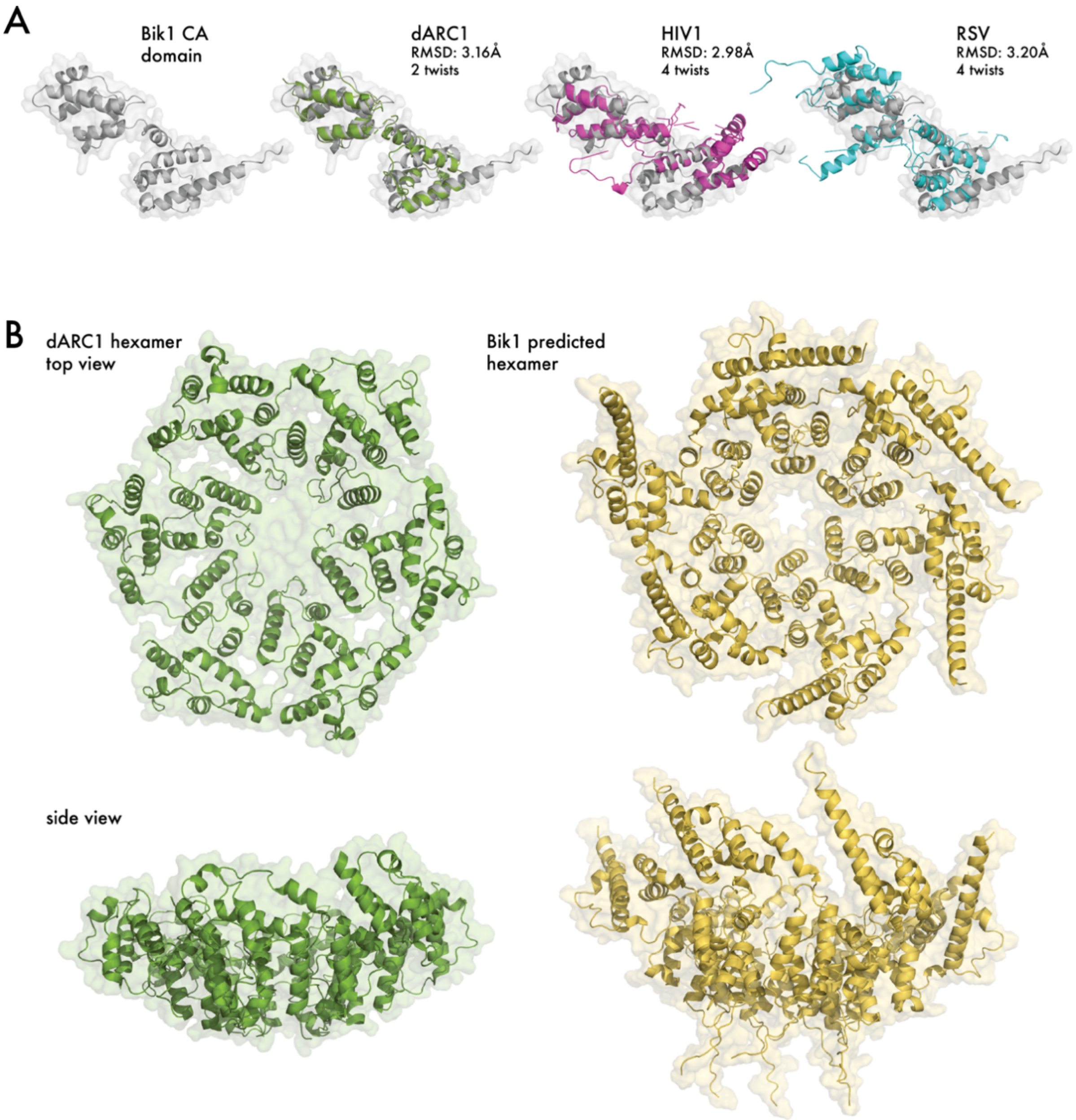


***Supplementary Figure 2. Bik-1 Gag contains a structurally conserved CA domain.***

*A) Structural alignments of the Bik1 CA domain to CA domains from D. melanogaster dARC1 (PDB: 6TAP-D (2)), HIV1 (PDB: 6OMT-A (3)), and Rous Sarcoma virus (PDB: 5A9E-A (4)). Root mean square deviation (RMSD) of alignments calculated using the flexible alignment solver, FATCAT 2.0; "twists" refers to number of added breaks in the chain being aligned to the template. B) Comparison of cryo-electron microscopy structure of dARC1 hexamer (PDB: 6TAS (2)) with AlphaFold2.0 predicted hexameric structure of Bik1 CA domain (2$^{nd}$ ranked), showing conserved cyclic C6 symmetry. Mature Gag capsids are formed from higher order combinations of hexamers and pentamers.*

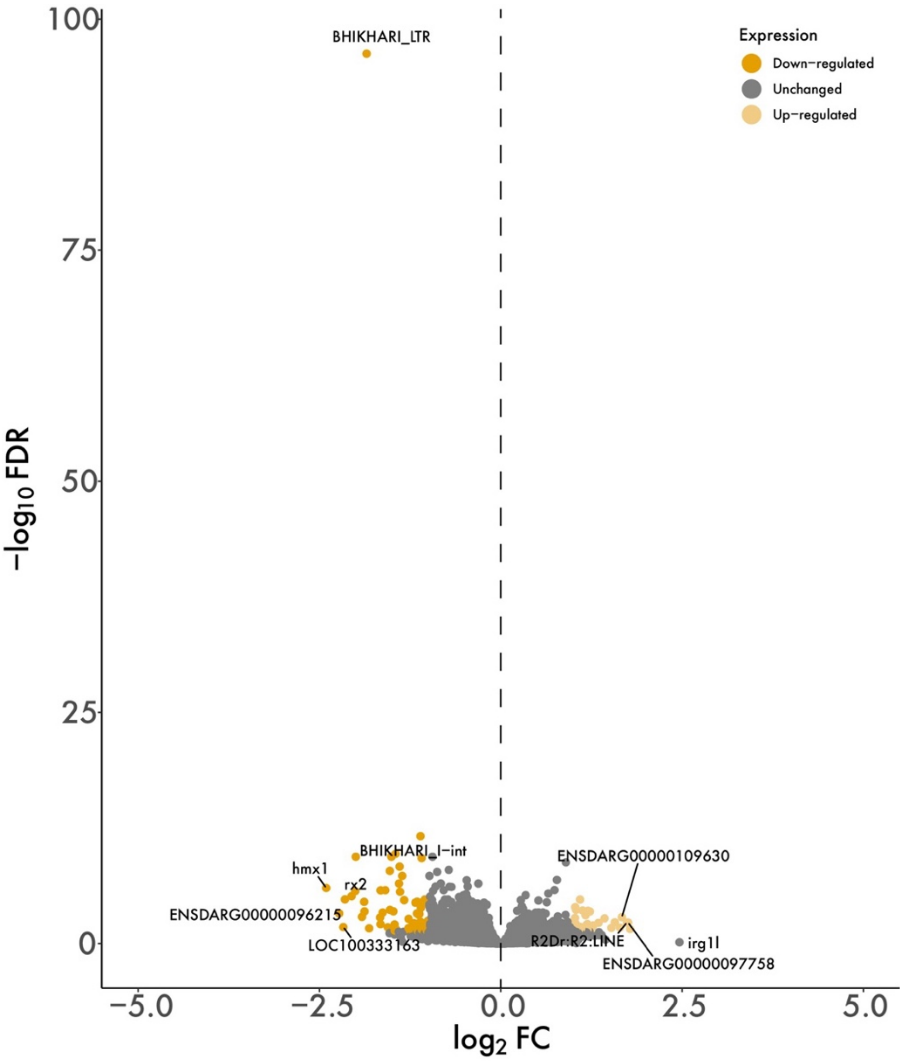


***Supplementary Figure 3. Bik-1-LNA depleted Bik-1***

*Volcano plot of RNA-seq from embryos injected with either 11pg Bik-1-LNA or scramble-LNA at one-cell stage, and collected at 8, 11, 14, 18 hpf. Time-points were treated as batch effects and regressed out. False discovery rate threshold 0.05, minimum fold-change 2.0.*

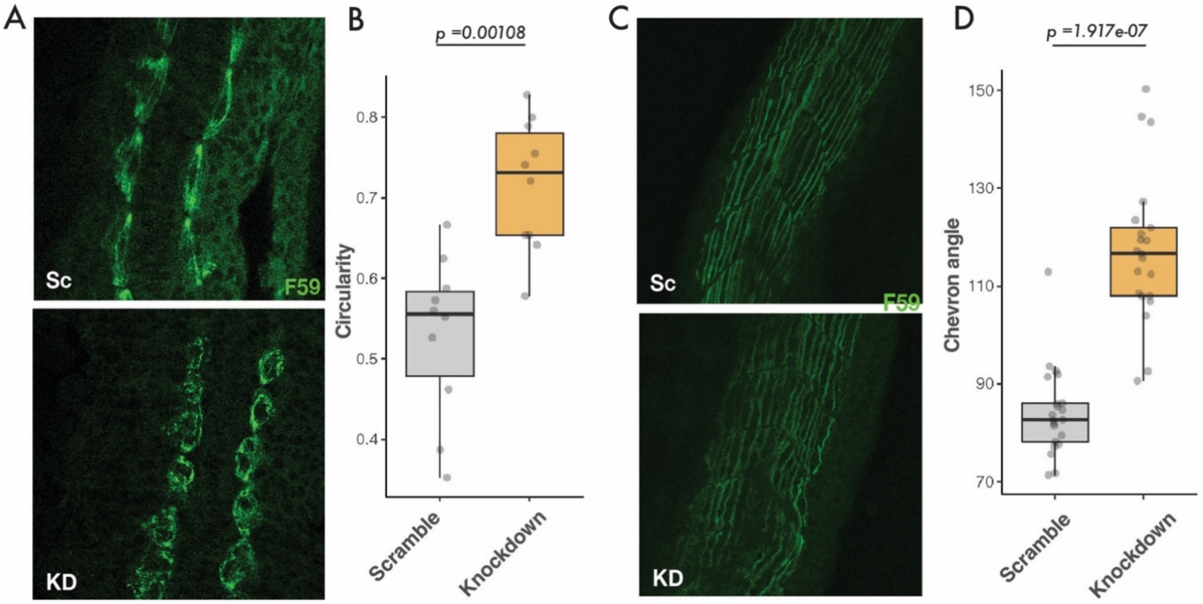


***Supplementary figure 4. Bik-1-LNA injected embryos shows defected phenotype on adaxial cells***
*A) Adaxial cells staining with F59 antibody at 12 hpf. B) Circularity quantification of adaxial cells at 12 hpf (Mann-Whitney Utest, p-value indicated in the figure). C) Slow muscle staining with F59 antibody at 16hpf. D) Measurement of the chevron angle on the slow muscles 16 hpf (Mann-Whitney U test, p-value indicated in the figure).*

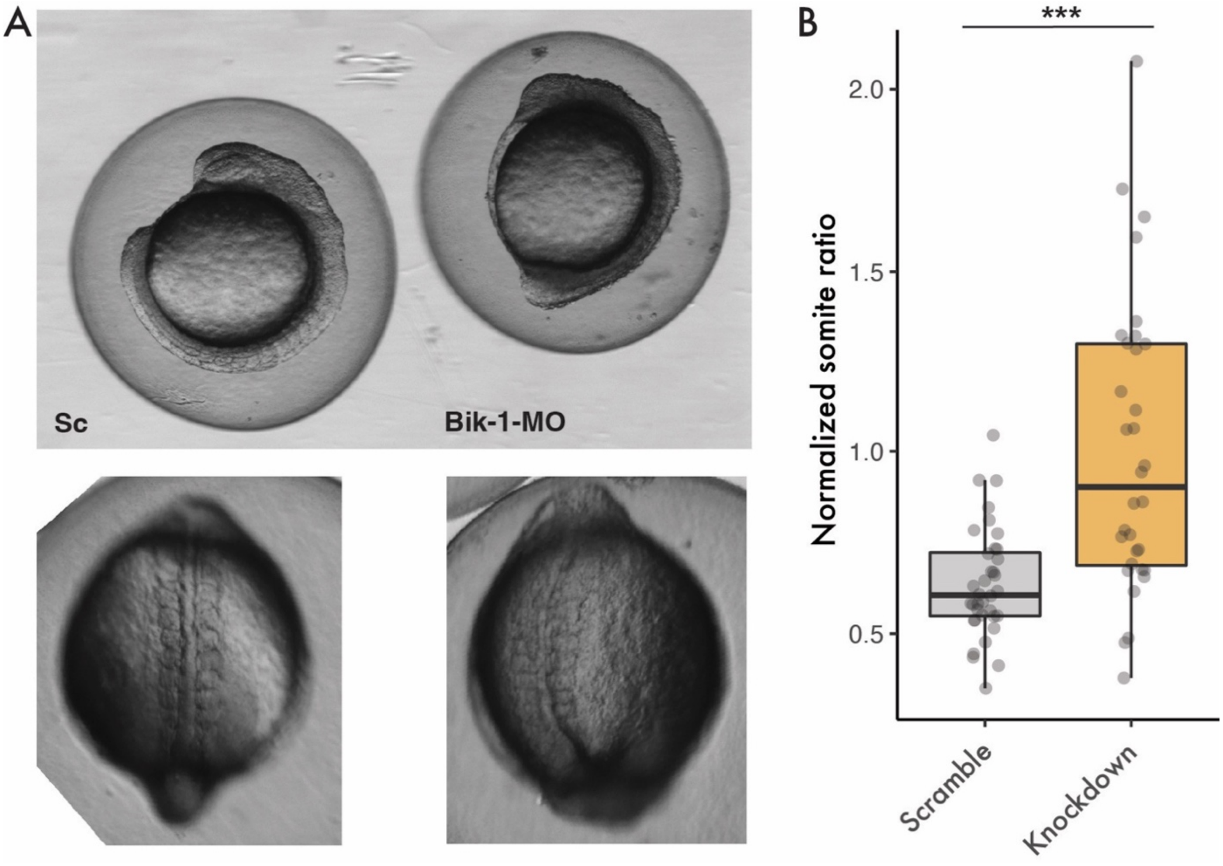

***Supplementary figure 5. Bik-1 MO injected embryos shows similar phenotype as Bik-1-LNA injected embryos***

*A) Embryos injected with either 2.7 ng/nl Bik-1 or scramble Morpholino and collected at 10-somite stage. 54 out of 59 scramble-MO injected embryos showed wildtype phenotype whereas 48 out of 60 Bik-1-MO showed the shortened axis and elongated somites phenotype. B) Normalized ratios of somite width/length across multiple injections and treatments (Mann-Whitney U test, p-value < 0.0001). SC, scramble-MO; Bik-1, Bik-1-MO.*

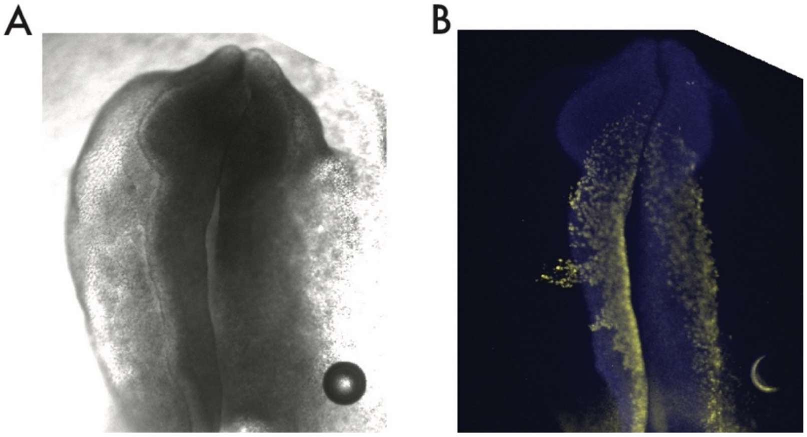


***Supplementary Figure 6. Neural tube defect in Tfap2α-Bik injected embryos***

*A) Tfap2α-mCherry (left) and Tfap2α-Bik (right) were injected and electroporated into chicken embryos. At HH9-0, Tfap2α-Bik injected side of embryos show neural tube defects but not the Tfap2α-mCherry side. Brightfield view of the embryo. B) Embryos were hybridized with Sox2 (blue) as a neural tube marker and Sox9 (yellow) as a neural crest cell marker.*

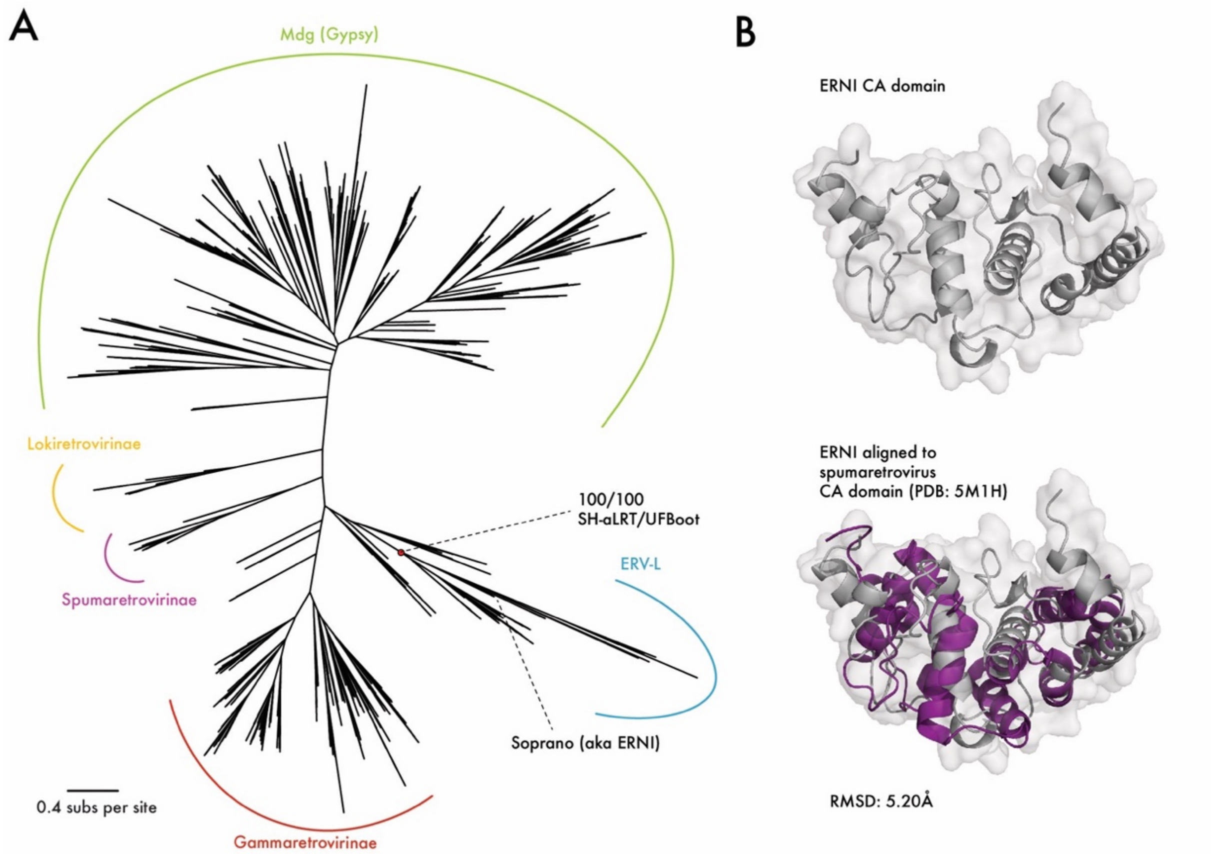


***Supplementary Figure 7. ERNI is a Gag capsid protein from an endogenous retrovirus***
*A) Phylogenetic tree of reverse transcriptase domain sequences from diverse retrovirus and LTR families. The reverse transcriptase domain of Soprano is confidently placed (Shimodaira–Hasegawa approximate likelihood ratio test > 0.85; ultra-fast bootstrap > 95) within a clade that is a sister to the ERV-L family of mammalian endogenous retroviruses. B) CEalign ((5)) structural alignment of ERNI CA domain with the CA domain of a spumaretrovirus Gag protein (PDB: 5M1H (6)).*

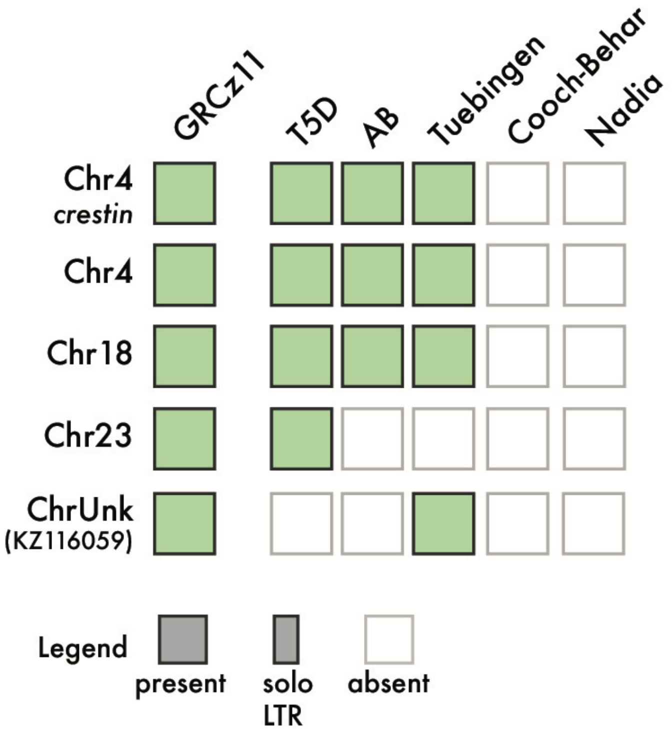


***Supplementary Figure 8. Bik-2 insertion polymorphisms***
*Analysis of presence/absence insertion polymorphisms from five Danio rerio genome assemblies for the five fully intact Bik-2 copies in the reference (GRCz11) genome.*

**Supplementary References**


1. J. Wang, G. Z. Han, *mBio* **13**, e0018722 (2022).
2. S. Erlendsson *et al.*, *Nat Neurosci* **23**, 172 (2020).
3. P. T. Huang *et al.*, *Cell Rep* **28**, 2373 (2019).
4. F. K. Schur, R. A. Dick, W. J. Hagen, V. M. Vogt, J. A. Briggs, *J Virol* **89**, 10294 (2015).
5. I. N. Shindyalov, P. E. Bourne, *Protein Eng* **11**, 739 (1998).
6. N. J. Ball *et al.*, *PLoS Pathog* **12**, e1005981 (2016).
