## Supplementary Data Legends for "Gag proteins encoded by endogenous retroviruses are required for zebrafish development"

**Supplementary Data Files**

**Supplementary Data 1.**
Aligned fasta files and corresponding phylogenetic trees from A) Bik-1 and B) Bik-2 insertions in *Danio rerio* reference genome (GCF_000002035.6_GRCz11).

**Supplementary Data 2.**
BAM files mapping full-length proviral Bik-1 and Bik-2 insertions in the *Danio rerio* reference genome to various other *D. rerio* genome assemblies, providing evidence for presence of polymorphic loci.

**Supplementary Data 3.**
Differential enrichment analysis of RNA-seq data from Bik-1 LNA knock-down experiment with time point differences regressed out.

**Supplementary Data 4.**
A) Aligned fasta file and corresponding phylogenetic tree of ENS-3 and related retroelement reverse transcriptase domains. B) Aligned fasta file and phylogenetic tree of ENS-1 insertions in the jungle fowl genome (GCF_000002315.6).
